# Inference of Rift Valley Fever pathogenesis in *Bos taurus* using a gene co-expression network

**DOI:** 10.1101/2021.11.28.470222

**Authors:** John K. Gitau, Rosaline W. Macharia, Kennedy W. Mwangi, Nehemiah M. Ongeso, Edwin Murungi

## Abstract

**Background:** Rift Valley Fever (RVF) is a viral disease caused by the Rift Valley Fever virus and spread mainly by the *Aedes* and *Culex* mosquito species. The disease primarily infects domestic animals such as sheep, goats, and cattle, resulting in a spectrum of clinical outcomes including morbidity, massive storm abortions, and neonatal fatalities. RVF outbreaks are closely linked to above-average rainfall and flooding, which create an ideal environment for mosquitos to breed, multiply, and transmit the virus to animals. The outcomes of human RVF infection range from self-limiting febrile illness to potentially fatal hemorrhagic diatheses and miscarriage in pregnant women. Collectively, the economic losses due to the zoonotic RVF disease is immense.

**Methods:** Using the Weighted Gene Co-expression Network Analysis (WGCNA) package, RNA-Seq data generated from five healthy *Bos taurus* steer calves aged 4-6 months was obtained from the Gene Expression Omnibus (GEO) database (Accession number GSE71417). The data was utilized to construct a gene co-expression network. Enriched modules containing genes potentially involved in RVF infection progression were identified. Moreover, using the Multiple Expectation Maximizations for Motif Elicitation (MEME) suite, consensus regulatory motifs of enriched gene clusters were deciphered and the most abundant putative regulatory motif in each enriched module unveiled by comparative analysis with publicly available motifs using the TOMTOM motif comparison tool. The potential roles of the identified regulatory motifs were inferred by literature mining.

**Results:** The constructed gene co-expression network revealed thirty-three (33) modules, nine of which were enriched for Gene Ontology terms linked to RVF pathogenesis. Functional enrichment in two (red and turquoise) of the nine modules was significant. ASH1-like histone lysine methyltransferase and Astrotactin were the hub genes for the red and turquoise modules respectively. ASH1-like histone lysine methyltransferase gene is involved in chromatin epigenetic modification while Astrotactin is a vertebrate-specific gene that plays an important role in neurodevelopment. Additionally, consensus regulatory motifs located on the 3' end of genes in each enriched module was identified.

**Conclusions:** In this study, we have developed a gene co-expression network that has aided in the unveiling of functionally related genes, intramodular hub genes, and immunity genes potentially involved in RVF pathogenesis. The discovery of functional genes with putative critical roles in the establishment of RVF infection establishment will contribute to the understanding of the molecular mechanism of RVF pathogenesis. Importantly, the putative regulatory motifs identified are plausible targets for RVF drug and vaccine development.

## Background

Rift Valley Fever (RVF) is a zoonotic disease caused by the Rift Valley Fever virus (RVFV), a triple segmented negative strand RNA virus [1], which is endemic throughout Africa [2]. The RVF primarily infects domestic animals such as sheep, goats, and cattle, resulting in a variety of clinical symptoms including massive storm abortions and numerous neonatal fatalities [3]. Although majority of humans infected with RVFV experience self-limiting febrile illness, 1-2% of infected people develop severe symptoms such as hemorrhagic diatheses and miscarriage in pregnant women. Mosquitoes act as vectors and reservoirs of RVFV [4] with the *Aedes* species enabling transovarian transmission while *Culex* and *Anopheles* species promote transmission to new hosts during routine blood feeding [3,5]. The RVF outbreaks are closely linked to above-average rainfall and flooding which offer an ideal environment for mosquito breeding and multiplication [1].

The depiction of interactions and molecular relationships using biological networks is important for the understanding of biological systems [6]. Biological systems include protein-protein interaction (PPI) networks that illustrate protein interactions in cellular systems, metabolic networks that depict metabolic fluxes, and gene regulatory networks that represent the interaction of different cellular modulators of gene expression and protein synthesis [7]. We have constructed a gene co-expression network, representing genes with similar expression patterns across multiple RNA-Seq samples [8]. In gene co-expression networks, nodes represent genes, while edges signify existence of co-expression relationships between genes [8,9]. Usually, genes with similar expression profiles across various samples are connected [6]. Co-expression networks aid in the understanding of system-level molecular interactions driving cellular processes. The modules (clusters) revealed in these networks generally contain genes with similar expression patterns that probably regulate analogous biological processes [6,9]. Moreover, modular genes tend to contain similar regulatory motifs, short nucleotide sequences that bind transcription factors to modulate gene expression [10]. Transcription factors have been shown to be tractable drug and vaccine targets for cancer and parasitic diseases [10, 11, 12].

In the present study, a *Bos taurus* gene co-expression network for the inference of RVF pathogenesis has been constructed from which, enriched modules have been identified and annotated. Moreover, immune response genes and putative consensus transcription factor binding motifs have been uncovered. The transcription factors binding to these motifs potentially play an important role in the pathogenesis of RVF and are thus plausible drug and vaccine targets.

## Materials and Methods

### Dataset retrieval and pre-processing

A total of 45 *Bos taurus* RNA-Seq data samples for a 21-day time course post vaccination with arMP-12 vaccine were obtained from the Gene Expression Omnibus (GEO) database [12] (Accession number GSE71417). The quality of the datasets was evaluated using FastQC version 0.11.7 (http://www.bioinformatics.babraham.ac.uk/projects/fastqc) prior to downstream analysis. To enable seamless analysis, quality reports obtained for each sample were combined into a single webpage using MultiQC version 1.6 [13]. Trimmomatic version 0.39 [14] was used to remove ambiguous sequences originating from adapters additions during Polymerase Chain Reaction (PCR), with the following parameters: ILLUMINACLIP: TruSeq3-SE:2:30:10 LEADING:3 TRAILING:3 SLIDINGWINDOW:4:15 MINLEN:36. After adapter removal, the quality of the samples was rechecked using FastQC version 0.11.7 (http://www.bioinformatics.babraham.ac.uk/projects/fastqc), and MultiQC version 1.6 [13].

### Reads alignment and filtering

Five of the forty-five (45) samples were controls, hence excluded from subsequent analyses. High quality reads from forty (40) remaining samples were mapped onto the *Bos taurus* reference genome using HISAT2 version 2.1.0 [15]. The *Bos taurus* reference genome (ARS-UCD1.2, INSDC Assembly GCA 002263795.2) obtained from the Ensembl database [16] was indexed prior to read alignment to accelerate the location of potential alignment sites and retrieval of genomic coordinates. The alignment obtained was converted to binary format and sorted to facilitate easier navigation using SAM tools version 1.9 [17]. Following sorting, the binary files were indexed, and a count matrix of reads uniquely mapping onto exons of a given gene generated by joining the single tables containing counts for all samples into a single table (data frame). All non-coding genes were expunged from the final count matrix. Additionally, low count genes were removed from the final count matrix using the filterByExpr function of the edgeR package version 3.8 [18]. The final count matrix was exported to R version 4.0 (https://www.r-project.org/) for further analysis.

### Construction and visualisation of a *Bos taurus* weighted gene co-expression network

The WGCNA version 1.69 R package was used to generate the *Bos taurus* gene co-expression network [19]. The first step involved log transformation of the discreet count matrix into a continuous distribution with a poisson-like distribution. This was followed by the construction of a similarity matrix which was subsequently converted into an adjacency matrix, using soft-thresholding power. Using WGCNA’s TOMsimilarity function, a Topological Overlap Matrix (TOM) was then computed from the adjacency matrix. The TOM was then converted into a dissimilarity matrix (1-TOM), which, in conjunction with average linkage hierarchical clustering, was used to group genes into modules based on their expression similarities. The flashClust function in WGCNA was used to generate a gene dendrogram (clustering tree) while the Dynamic Tree Cut function was used to decipher gene modules (clusters). Hub genes for each module were unveiled using the chooseTopHubInEachModule function. The interaction in the constructed *Bos taurus* gene co-expression network was visualised using Cystoscape version 3.8.0 [20], with the various gene modules being depicted in different colours.

### Gene ontology and functional enrichment analysis

The modules (gene clusters) identified were analysed for enrichment of gene ontology (GO) terms using the GOSEQ version 1.36.0 R package [21], and thereafter, redundancy in the long GO terms minimised using REVIGO (Reduce and Visualize Gene Ontology) [22]. A two-dimensional scatter plot was used to visualise the most informative and biologically relevant non-redundant GO terms in each enriched module.

### 3’ UTR regulatory motif prediction and abundance analysis

The Multiple Expectation Maximizations for Motif Elicitation (MEME) suite [23] was used for the *de novo* identification of motifs with potential gene regulatory roles on the 3’ end of genes in all enriched modules. The DNA sequences of all genes in the nine enriched modules were obtained from the BioMart database [24]. All duplicated gene sequences, genes lacking 3’ sequences, and sequences containing fewer than eight nucleotides were excluded from the input data for MEME. To determine motif similarity, the most abundant putative regulatory motif in each module was compared to motifs in public databases using the TOMTOM motif comparison tool. The significance and potential roles of the identified regulatory motifs was inferred from published literature.

## Results

### *Bos taurus* RNA-Seq read mapping and count matrix generation

After quality evaluation, the 40 high-quality samples were mapped onto the *Bos taurus* reference genome (ARS-UCD1.2, INSDC Assembly GCA 002263795.2) with an average mapping efficiency of 83%. Count matrices were constructed for each gene (genes as rows and samples as columns) (Additional File 2). Non-protein coding biotypes such as rRNA, lncRNA, snRNA, miRNA, snoRNA, miRNA, Pseudogenes, misc RNA, and processed pseudogenes were excised from each count matrix. Additionally, 5,627 non-coding genes and 9232 genes with low counts reads across the 40 samples were removed. The final count matrix contained 12411 genes.

### *Bos taurus* weighted gene co-expression network

All the 12,411genes in the final count matrix were log transformed and used to construct a gene co-expression network based on WGCNA analysis. The count matrix was converted into a similarity matrix by combining the benefits of Pearson correlation and Euclidean distance. Thereafter, saturation analysis (Figure 1) performed revealed that a soft thresholding power of 7 with a scale-free topology fitting index (R^2^) of ≥ 0.8 was optimal for converting the similarity matrix to an adjacency matrix. From the adjacency matrix, Dynamic Tree Cut function and hierarchical clustering generate 33 distinct modules (Figure 2). The 57 genes in the gray module were not assigned to any of the other modules and were thus excluded from further analysis because their expression patterns did not match those of existing modules. The most connected genes in each module (hub genes) were identified as shown in Additional File 3. containing 2314 genes, the turquoise module had the most genes while the darkolivegreen module with 37 genes had the least genes. The interaction of the various modules and the genes therein was determined and illustrated in Figure 3.

**Fig 1:**
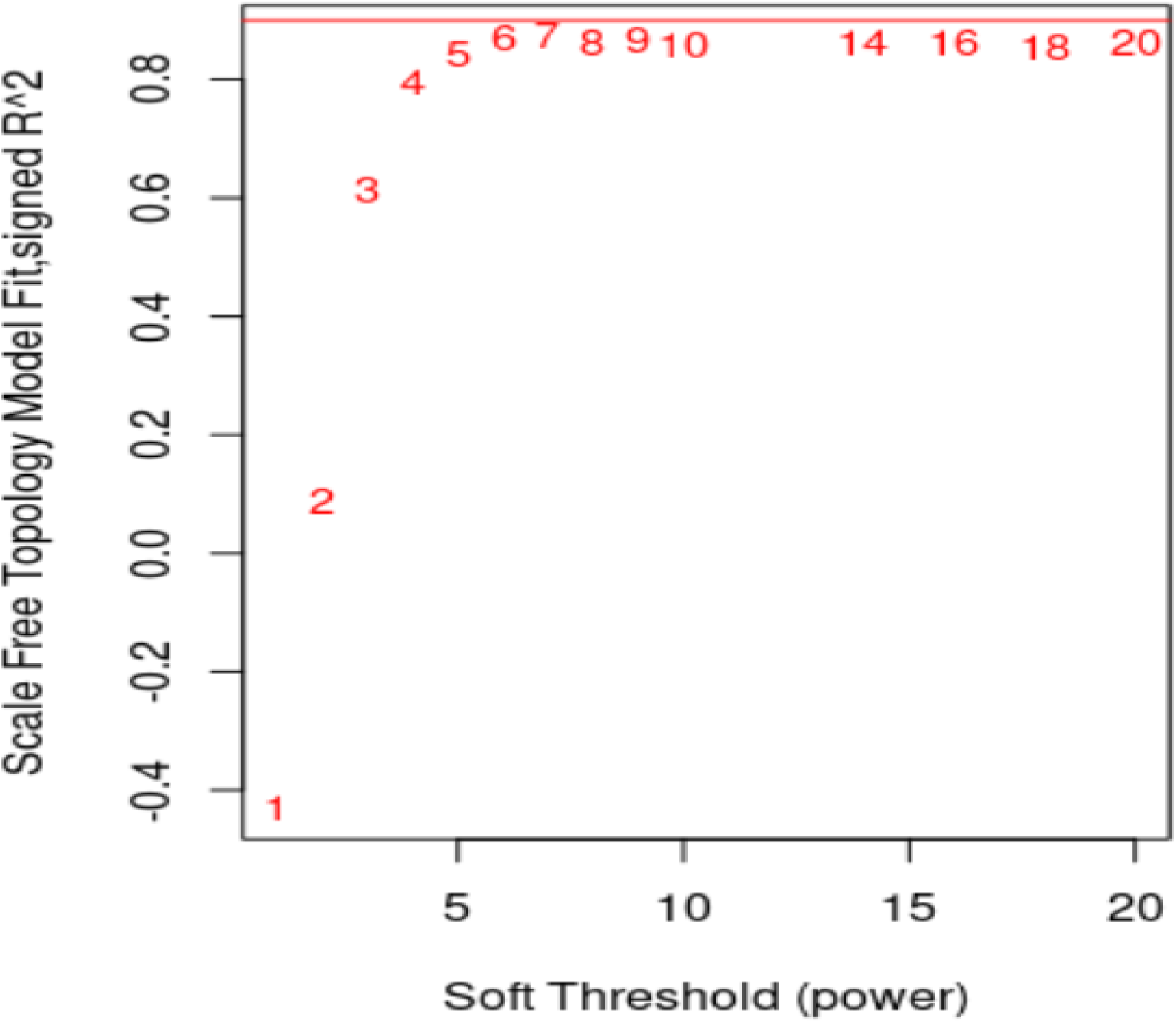
A saturation curve for determining the appropriate soft thresholding power for transforming the similarity matrix of 40 *Bos taurus* samples into an adjacency matrix. The x-axis depicts various power values, while the y-axis depicts the cut-off values used to select an appropriate power value. Using a cut-off of 0.96, the best value was determined to be 7, the point at which the curve begins to plateau.

**Fig 2:**
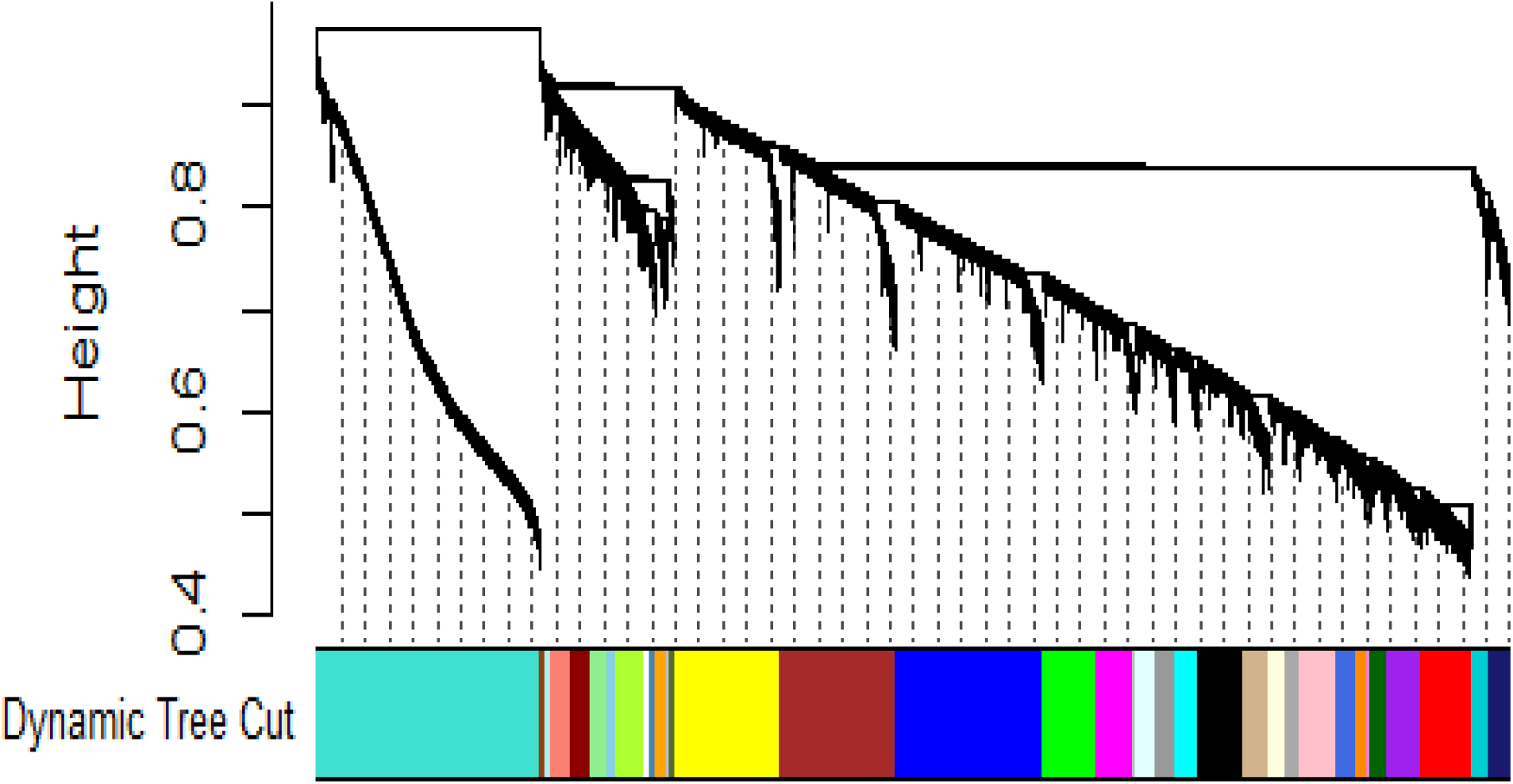
A color-coded plot of the 33 gene modules of the *Bos taurus* gene co-expression network (x-axis). The WGCNA R package’s dynamic tree cut function cut the gene dendrogram at different heights based on expression similarities (y-axis), resulting in the different module colors.

**Fig 3:**
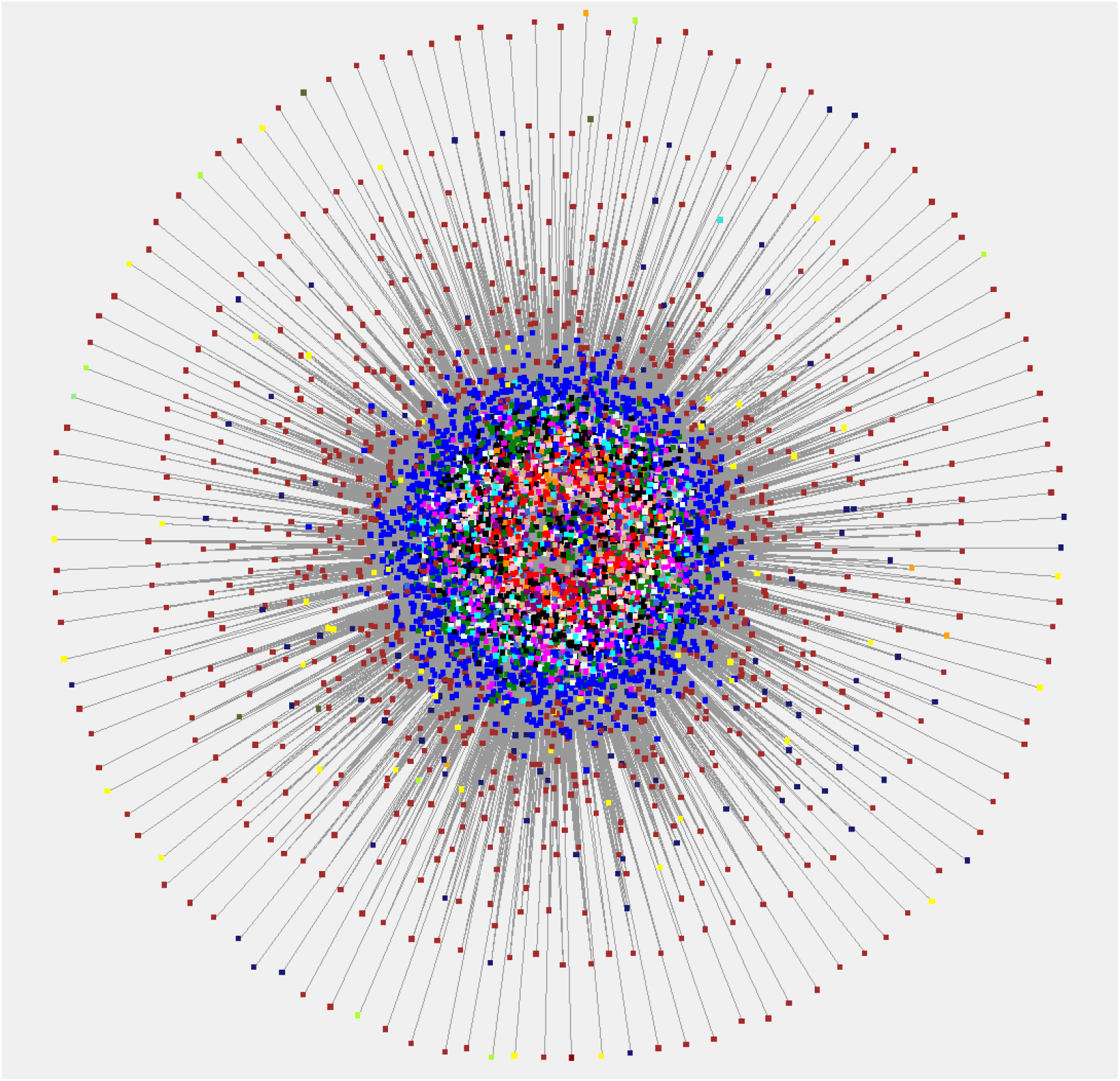
An interaction network depicting the interactions between genes in the 33 distinct modules of the *Bos taurus* gene co-expression network. Each module is depicted in a different colour to illustrate the interactions between different modules.

### Identification of hub genes in enriched modules

Table 1 shows the hub genes for the nine enriched modules with over-represented GO terms.

**Table 1.** Hub genes in the nine enriched modules. For each module, the module’s colour, the number of genes, the hub gene name and the hub gene Ensembl accession are shown.

| **Module color** | **Number of genes** | **Hub gene name** | **Hub gene symbol** |
| --- | --- | --- | --- |
| Cyan | 233 | DExH-box helicase 9 | DHX9 |
| Green | 561 | integrin subunit beta 1 | ITGB1 |
| Light green | 194 | Integral membrane protein 2B | ITM2B |
| Orange | 121 | ornithine decarboxylase antizyme 1 | OAZ1 |
| Red | 538 | ASH1 like histone lysine methyl transferase | ASH1L |
| Steel blue | 56 | Ribosomal protein S18 | RPS18 |
| Tan | 257 | Protein kinase C beta | PRKCB |
| Turquoise | 2314 | Astrotactin 2 | ASTN2 |
| White | 79 | Ribosomal protein L23a | RPL23A |

### Gene ontology and functional enrichment analysis

Gene Ontology (GO) analysis revealed that nine out of thirty-three (33) modules (cyan, green, light green, orange, red, steel blue, tan, turquoise and white) were enriched for GO terms. The red and turquoise modules were significantly enriched for GO terms related to RVF pathogenesis. The red module was enriched for terms such as calcineurin-NFAT signaling cascade, protein ubiquitination, and ubiquitin-protein ligase activity while the turquoise module was enriched for terms like dendrite, G protein-coupled receptor activity, response to stimuli, and xenobiotic metabolic process. Filtering of the GO terms in the nine enriched modules for redundancy using Revigo, which uses a semantic similarity approach, resulted in two-dimensional scatter plots. The plot for the turquoise module is shown in Figure 4.

**Fig 4:**
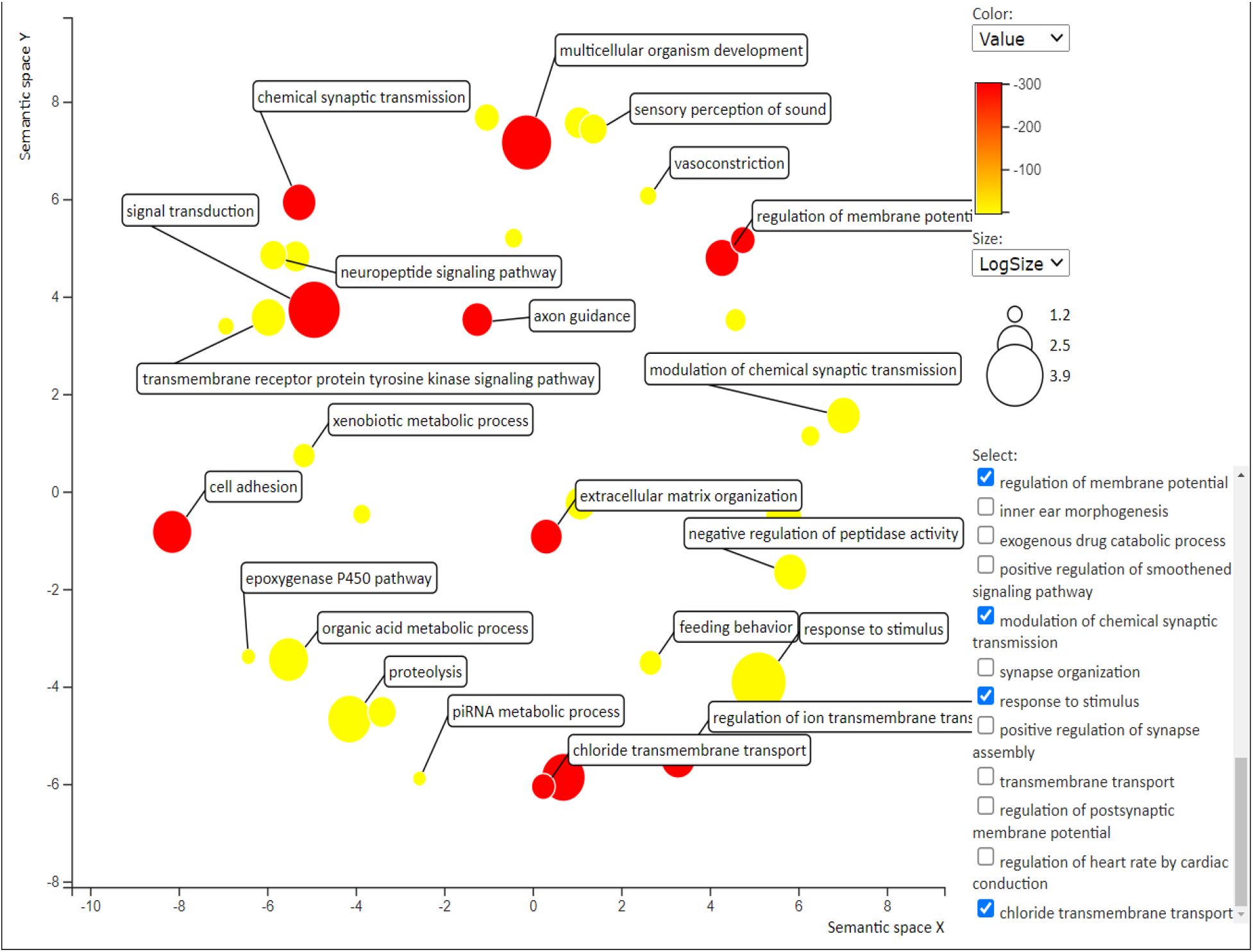
A two-dimensional scatter plot of non-redundant biological process Gene Ontology (GO) terms in the turquoise module. After redundancy reduction, the GO terms displayed are the most significant. Different bubble colors indicate different P-values used (legend in upper right corner), while bubble sizes indicate the frequency of GO terms from source databases.

### Identification and abundance analysis of regulatory motifs

The most abundant motif in each enriched module was identified and tabulated (table 2). The most abundant motif was uncovered by ascertaining whether it is present in the majority of the genes which putatively indicates its importance in the functioning of the module.

**Table 2.**
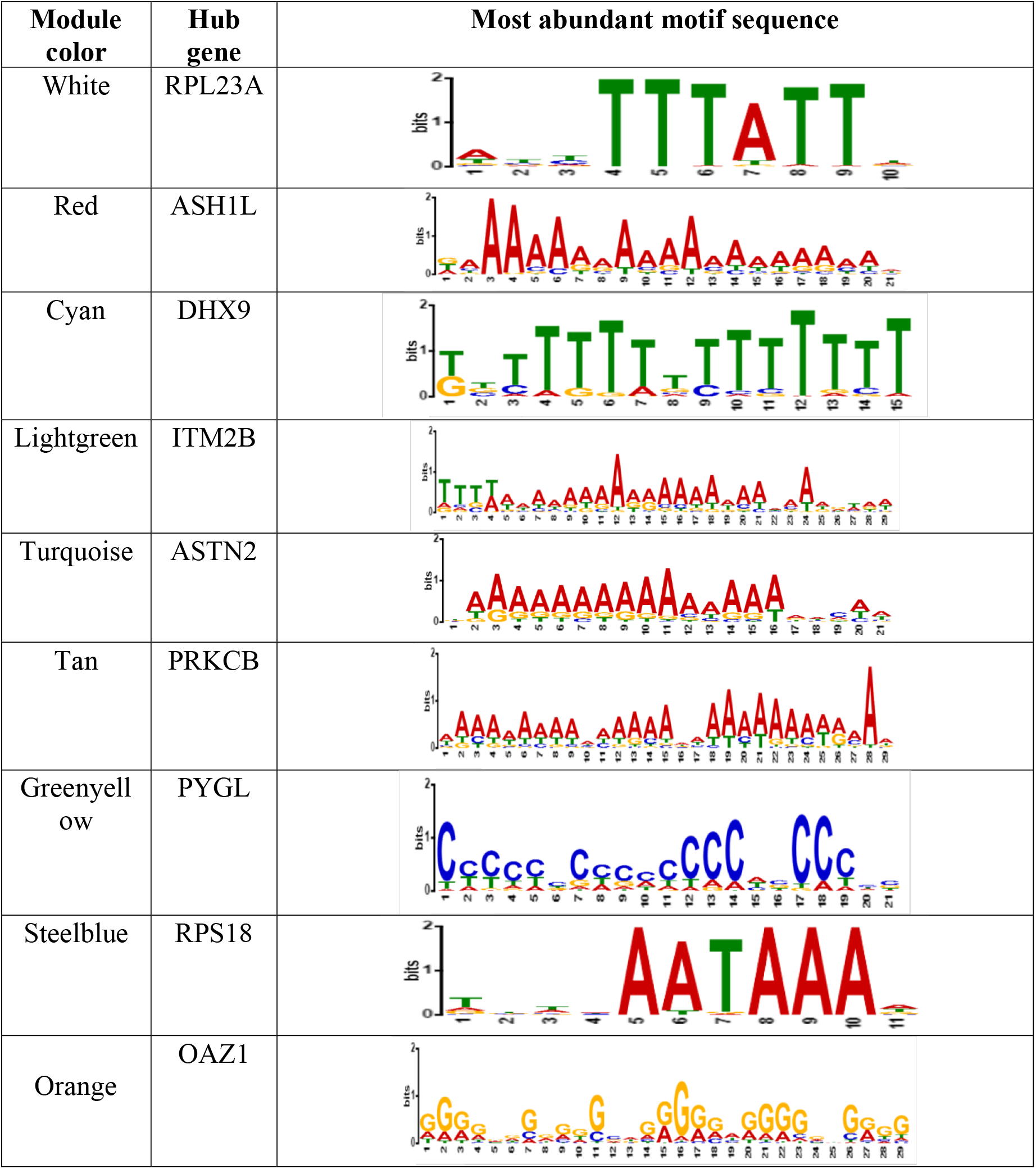
A list of the nine (9) enriched modules, their hub genes, and the most abundant regulatory motif sequences.

## Discussion

Using the WGCNA methodology, a *Bos taurus* gene co-expression network was constructed in which thirty-three (33) modules (Figure 2) with similar expression patterns were deciphered. For each enriched module (Table 1), hub genes (the most connected genes) which plausibly perform important biological functions due to their extensive connectivity and may thus serve as representatives of their respective modules, were identified. By performing gene ontology (GO) and functional enrichment analysis in GOSEQ [21], enriched modules and immune genes associated with RVF infection were uncovered. The functional roles attributed to the enriched modules include stimulus response, signaling and transduction, xenobiotic metabolic process, and protein ubiquitination. In nine enriched (biologically significant) modules, consensus regulatory motifs that may aid in the determination of potential gene regulatory mechanisms were identified using MEME [23] identified consensus regulatory motifs in nine enriched (biologically significant) modules to uncover their potential regulatory mechanisms.

It has been demonstrated that genes with similar expression patterns are likely to have related functions, be components of analogous cellular regulatory and signaling processes [9,25]. Majority of GO terms associated with RVF pathogenesis were significantly enriched in the red and turquoise modules in the constructed network. The hub genes for the red and turquoise modules were ASH1-like histone lysine methyltransferase and Astrotactin 2. ASH-1 like histone lysine methyltransferase, is involved in chromatin epigenetic modification and is associated with the transcribed region of developmentally important genes such as the homeobox (HOX) gene [26]. Repression of the HOX gene by ASHI-like histone lysine methyltransferase may be responsible for the fetal malformations seen in RVF infection. It has been shown that bovine oocytes and early embryos express HOX genes for vertebrate body axis patterning beginning with gastrulation [27]. Similar to other viruses, RVF replication is through cellular epigenetic processes encoded by ASH1-like histone lysine methyltransferase. Therefore, the interaction between ASH1-like histone lysine methyltransferase and the HOX gene may be disrupted in RVF. ASHI-1 has been shown to bind to the HOX loci in K562 cells and ASH-1 knockdown regulates HOX gene expression [28]. This observation could explain the observed fetal malformation and storm abortions in RVF-infected pregnant cattle. Since the ASH-1 and HOX gene association may be compromised following RVFV establishment and subsequent replication, a *Bos taurus* fetus normal body patterning is disrupted and may lead to abortion if the disruption is severe. Because ASH1, like histone lysine methyltransferase, is involved in epigenetic chromatin modification, evidence suggests that a change in chromatin and chromatin-modifying enzymes results in a persistent dysregulation of metabolic phenotype, thereby amplifying disease pathogenesis [29]. Astrotactin 2, a perforin-like integral membrane protein, is a vertebrate-specific gene that plays an important role in neurodevelopment [30]. Perforins are released natural killer cells (cytotoxic T lymphocytes) and target virus-infected and host transformed cells [30].

Gene ontology and functional enrichment analysis determine whether a gene co-expression module has a biological meaning, such as a metabolic pathway, or if it is noise caused by tissue contamination, false positives, or artifacts [29]. In this study, nine modules were identified as being enriched for RVF pathogenesis-related terms. Ontology terms associated with ribosomal structure and function were enriched for genes in the white, Steelblue, and Greenyellow modules. Enriched ontology terms included translation and cytosolic large ribosomal subunit. The RVFV like other viruses relies on the host protein synthesis machinery to translate its mRNA and synthesise proteins required for infection progression [31]. The interferon-inducible protein kinase R (PKR), encoded by the PRKRA gene, is present in the green module and plays an important role in host innate immunity by limiting viral particle replication and propagation [32]. The ribosome plays an important role in the translation of interferon-inducible protein kinase R, which is involved in bovine immune responses. The S-segment of the viral nucleic material encodes the nonstructural (NS) protein, a major virulence factor, upon Rift Valley Fever virus invasion, promoting TFIIH p62 degradation [5].

The interferon regulatory factor 1 (IRF1) gene is a key gene present the enriched Lightgreen module. Interferon-mediated signaling pathway and biosynthetic process are two gene ontology terms associated with this gene. The IRFI promotes host defense mechanisms by modulating autoimmunity, as well as innate and adaptive immune responses [10]. *Bos taurus* expresses this gene in response to RVF viral infection in order to mount an effective response.

Terms like calcineurin-NFAT signaling cascade and protein ubiquitination were enriched within the red module. The calcineurin/nuclear factor of activated T cells (NFATs) signaling pathway is involved in T cell-mediated adaptive immune responses as well as myeloid cell regulation of the innate immune response [33]. The *Bos taurus* immune system counters RVFV infection by releasing macrophages and activated T cells to destroy the virus. It has been reported that ubiquitination is an important signal transduction mechanism [34]. Ubiquitination promotes T cell activation and differentiation, allowing a host to appropriately respond, following pathogen such as RVF virus invasion [34]. Ubiquitination regulates pattern recognition receptor signaling, allowing it to mediate both innate immune response and dendritic cell maturation. Pattern recognition receptor signaling initiates an adaptive immune response. Macrophages are innate immune cells that express distinct families of pattern recognition receptors (PRRs) that mediate phagocytosis and signal transduction by recognising pathogen-associated molecular patterns (PAMPs) [35]. The interaction between macrophages and specific pathogens promotes the production of antimicrobial factors and inflammatory mediators, resulting in the pathogen’s destruction [34].

Monocytes and macrophages by expressing cytokines and chemokines play a key role in the innate immune response [36]. *Bos taurus* produces monocytes and macrophages in response to RVF virus infection in order to destroy the RVF viral particles. Macrophages express a number of G protein-coupled receptors (GPCRs), which play important roles in cell differentiation and cell function regulation [37]. The GPCRs which are involved in the regulation of immune cell response, are critical therapeutic targets in a variety of diseases including diabetes [37]. The presence of GPCRs in the turquoise module indicates the importance of GPCR-linked pathways in the pathogenesis of RVF and suggests that activators of bovine GPCRs could be used as RVF therapeutics.

The most abundant motif in each enriched module is represented by the identified consensus regulatory motifs (Table 2). Because the consensus motif is present in the DNA sequence of the majority of genes in the respective module, the mechanism of regulation of these motifs may have an impact on the overall regulation of module activity.

## Conclusions

The gene co-expression network approach is a powerful approach for uncovering interactions between a virus and its host as demonstrated in the identification of *Bos taurus* functional genes in RVF infection. The identification of significantly enriched *Bos taurus* gene clusters in RVF infection will aid in the elucidation of the molecular mechanisms underlying RVF pathogenesis and potentially uncover novel tractable drug targets for RVF.

## Supporting information

Metadata

Over-represented and under-represented GO terms in nine enriched modules

non-redundant biological processes GO terms in the turquoise module

non-redundant molecular function GO terms in the turquoise module

non-redundant cellular compartment GO terms in the turquoise module

non-redundant biological function GO terms in the red module

non-redundant molecular function GO terms in the red module

non-redundant Cellular compartment Gene Ontology terms in the red module

non-redundant biological process Gene Ontology terms in the white module

non-redundant Molecular function GO terms in the white module

non-redundant cellular compartment GO terms in the white module

non-redundant biological activity GO terms in the Steelblue module

non-redundant Molecular function GO terms in the Steelblue module

non-redundant Cellular compartment GO terms in the Steelblue module

non-redundant cellular compartment GO terms in the green module

non-redundant biological activity GO terms in the Lightgreen module

non-redundant cellular compartment GO terms in the orange module

non-redundant cellular compartment Gene Ontology terms in the tan module

Cellular compartment GO terms in the cyan module

## Additional files

**Additional file 1**: The pipeline used to process the data and generate conclusions in this analysis is achieved at https://github.com/Jkgitau/RVF_WGCNA.

**Additional file 2: Table S1**. Samples metadata utilized in this study

**Additional file 3: Figure S2.** Scale free topology plot for selecting the power β for the signed correlation network. Scale free topology index (*y* axis) as a function of powers, β, 1 to 20 (*x* axis).

**Additional file 4: Table S2**. Over-represented and under-represented GO terms in nine enriched modules. The number of genes in each module, biological Process GO terms, Molecular Function GO terms and Cellular compartment GO terms are outlined.

**Additional file 5: Table S3.** A summary of non-redundant biological processes GO terms in the turquoise module.

**Additional file 6: Table S4.** A summary of the non-redundant molecular function GO terms in the turquoise module.

**Additional file 7: Table S5.** A summary of the non-redundant cellular compartment GO terms in the turquoise module.

**Additional file 8: Table S6.** A summary of the non-redundant biological function GO terms in the red module.

**Additional file 9: Table S7.** A summary of the non-redundant molecular function GO terms in the red module.

**Additional file 10: Table S8.** A summary of non-redundant Cellular compartment Gene Ontology terms in the red module.

**Additional file 11: Table S9.** A summary of non-redundant biological process Gene Ontology terms in the white module.

**Additional file 12: Table S10.** A summary of non-redundant Molecular function GO terms in the white module.

**Additional file 13: Table S11.** A summary of non-redundant cellular compartment GO terms in the white module.

**Additional file 14: Table S12.** A summary of non-redundant biological activity GO terms in the Steelblue module.

**Additional file 15: Table S13.** A summary of non-redundant Molecular function GO terms in the Steelblue module.

**Additional file 16: Table S14.**A summary of non-redundant Cellular compartment GO terms in the Steelblue module.

**Additional file 17: Table S15.** A summary of non-redundant cellular compartment GO terms in the green module.

**Additional file 18: Table S16.** A summary of non-redundant biological activity GO terms in the Lightgreen module.

**Additional file 19: Table S17.** A summary of non-redundant cellular compartment GO terms in the orange module.

**Additional file 20: Table S18.** A summary of non-redundant cellular compartment Gene Ontology terms in the tan module.

**Additional file 21: Table S19.** A summary of non-redundant Cellular compartment GO terms in the cyan module.

## Abbreviations

RVF: Rift Valley Fever
RVFV: Rift Valley Fever Virus
PPI: Protein-Protein Interactions
WGCNA: Weighted Gene Co-Expression Network Analysis
GPCR: G-protein coupled receptor
GO: Gene Ontology
MEME: Multiple Expectation Maximizations for Motif Elicitation
TOM: Topological overlap matrix

## Acknowledgements

The authors are thankful for the assistance accorded by Dr. Francis Makokha, head of research and innovation, Mt. Kenya University for his thoughtful insights while conducting the study. We acknowledge the University of Nairobi, Biochemistry department for providing a learning environment to complete this study.

## Ethics approval and consent to participate

Not applicable

## Consent for publication

Not applicable

## Availability of data and materials

The datasets supporting the conclusions of this study are included within the article and in the additional data files.

## Competing interests

The authors declare that they have no competing interests.

## Funding

Not applicable

## Author contributions

JKG, RWM, EM conceived experimental designs. JKG conducted the experiments, RWM, EM, KWM contributed materials/analysis tools. JKG, RWM, EM, KWM, NMO analyzed data. JKG developed the first manuscript draft. All authors read and approved the final manuscript.

## Author information

^1^University of Nairobi, Biochemistry department, P.O Box 30197 – 00100, Nairobi, Kenya. ^2^Kisii University, Department of Medical Biochemistry, P.O Box 408 – 40200, Kisii, Kenya, ^3^Jomo Kenyatta University of Agriculture and Technology, P.O Box 62000 – 00200, Nairobi, Kenya.

## Notes

https://www.ncbi.nlm.nih.gov/geo/query/acc.cgi?acc=GSE71417

### Competing Interest Statement

The authors have declared no competing interest.

