## Supplementary material for "Inference of Rift Valley Fever pathogenesis in *Bos taurus* using a gene co-expression network": Metadata

**Additional file 2: Table S1**. Samples metadata utilized in this study

| Run | Individual | Serologic_response_Status | Time |
|---|---|---|---|
| SRR2131563 | Animal 86 | vaccinated, not protected | Day 0 |
| SRR2131564 | Animal 88 | vaccinated, not protected | Day 0 |
| SRR2131565 | Animal 75 | vaccinated, not protected | Day 2 |
| SRR2131566 | Animal 86 | vaccinated, not protected | Day 2 |
| SRR2131567 | Animal 88 | vaccinated, not protected | Day 2 |
| SRR2131568 | Animal 74 | vaccinated, not protected | Day 3 |
| SRR2131569 | Animal 75 | vaccinated, not protected | Day 3 |
| SRR2131570 | Animal 86 | vaccinated, not protected | Day 3 |
| SRR2131571 | Animal 88 | vaccinated, not protected | Day 3 |
| SRR2131572 | Animal 74 | vaccinated, not protected | Day 4 |
| SRR2131573 | Animal 75 | vaccinated, not protected | Day 4 |
| SRR2131574 | Animal 86 | vaccinated, not protected | Day 4 |
| SRR2131575 | Animal 75 | vaccinated, not protected | Day 5 |
| SRR2131576 | Animal 76 | vaccinated, not protected | Day 5 |
| SRR2131577 | Animal 86 | vaccinated, not protected | Day 5 |
| SRR2131578 | Animal 88 | vaccinated, not protected | Day 5 |
| SRR2131579 | Animal 74 | vaccinated, not protected | Day 6 |
| SRR2131580 | Animal 75 | vaccinated, not protected | Day 6 |
| SRR2131581 | Animal 76 | vaccinated, not protected | Day 6 |
| SRR2131582 | Animal 86 | vaccinated, not protected | Day 6 |
| SRR2131583 | Animal 88 | vaccinated, not protected | Day 6 |
| SRR2131584 | Animal 74 | vaccinated, not protected | Day 7 |
| SRR2131585 | Animal 75 | vaccinated, not protected | Day 7 |

| | | | |
|---|---|---|---|
| SRR2131586 | Animal 76 | vaccinated, not protected | Day 7 |
| SRR2131587 | Animal 86 | vaccinated, protected | Day 7 |
| SRR2131588 | Animal 74 | vaccinated, not protected | Day 10 |
| SRR2131589 | Animal 75 | vaccinated, protected | Day 10 |
| SRR2131590 | Animal 76 | vaccinated, not protected | Day 10 |
| SRR2131591 | Animal 86 | vaccinated, not protected | Day 10 |
| SRR2131592 | Animal 88 | vaccinated, protected | Day 10 |
| SRR2131593 | Animal 74 | vaccinated, protected | Day 14 |
| SRR2131594 | Animal 75 | vaccinated, protected | Day 14 |
| SRR2131595 | Animal 76 | vaccinated, protected | Day 14 |
| SRR2131596 | Animal 86 | vaccinated, protected | Day 14 |
| SRR2131597 | Animal 88 | vaccinated, protected | Day 14 |
| SRR2131598 | Animal 74 | vaccinated, protected | Day 21 |
| SRR2131599 | Animal 75 | vaccinated, protected | Day 21 |
| SRR2131600 | Animal 76 | vaccinated, protected | Day 21 |
| SRR2131601 | Animal 86 | vaccinated, protected | Day 21 |
| SRR2131602 | Animal 88 | vaccinated, protected | Day 21 |
