## Supplementary material for "Inference of Rift Valley Fever pathogenesis in *Bos taurus* using a gene co-expression network": Over-represented and under-represented GO terms in nine enriched modules

**Additional file 4: Table S2**. Over-represented and under-represented GO terms in nine enriched modules. The number of genes in each module, biological Process GO terms, Molecular Function GO terms and Cellular compartment GO terms are outlined.

**Steelblue module (56 genes)**

Over-represented GO terms:

| category | num_in_subset | num_total | adj_pval | term | ontology |
|---|---|---|---|---|---|
| GO:0003735 | 17 | 104 | 0e+00 | molecular_function | structural constituent of ribosome |
| GO:0005840 | 14 | 99 | 0e+00 | cellular_component | ribosome |
| GO:0006412 | 14 | 164 | 5e-07 | biological_process | translation |

**White module (34 genes)**

**Over-represented GO terms:**

| category | num_in_subset | num_total | adj_pval | term | ontology |
|---|---|---|---|---|---|
| GO:0005840 | 17 | 99 | 0.0000002 | cellular_component | ribosome |

| | | | | | |
|---|---|---|---|---|---|
| GO:0003735 | 17 | 104 | 0.0000002 | molecular_function | structural constituent of ribosome |
| GO:0006412 | 17 | 164 | 0.0000006 | biological_process | translation |
| GO:0022627 | 6 | 24 | 0.0319196 | cellular_component | cytosolic small ribosomal subunit |

**Red module (378 genes)**

Over-represented GO terms:

| category | num_in_subset | num_total | adj_pval | term | ontology |
|---|---|---|---|---|---|
| GO:0005634 | 234 | 3117 | 0.0000000 | cellular_component | nucleus |
| GO:0005654 | 147 | 2064 | 0.0000789 | cellular_component | nucleoplasm |
| GO:0016567 | 39 | 311 | 0.0000789 | biological_process | protein ubiquitination |
| GO:0004842 | 27 | 179 | 0.0011837 | molecular_function | ubiquitin-protein transferase activity |
| GO:0061630 | 26 | 191 | 0.0015220 | molecular_function | ubiquitin protein ligase activity |
| GO:0003677 | 64 | 698 | 0.0021103 | molecular_function | DNA binding |
| GO:0033173 | 6 | 10 | 0.0030222 | biological_process | calcineurin-NFAT signaling cascade |

| | | | | | |
|---|---|---|---|---|---|
| GO:0005829 | 140 | 2063 | 0.0107478 | cellular_component | cytosol |
| GO:0006511 | 24 | 167 | 0.0107478 | biological_process | ubiquitin-dependent protein catabolic process |

Under-represented GO terms:

| category | num_in_subset | num_total | adj_pval | term | ontology |
|---|---|---|---|---|---|
| GO:0016021 | 71 | 2615 | 0.0029264 | cellular_component | integral component of membrane |
| GO:0005576 | 1 | 327 | 0.0076568 | cellular_component | extracellular region |
| GO:0005615 | 2 | 384 | 0.0133871 | cellular_component | extracellular space |
| GO:0016020 | 104 | 3274 | 0.0159715 | cellular_component | membrane |

**Cyan module (160 genes)**

Under-represented GO terms:

| category | num_in_subset | num_total | adj_pval | term | ontology |
|---|---|---|---|---|---|
| GO:0016021 | 21 | 2615 | 0.0046178 | cellular_component | integral component of membrane |
| GO:0016020 | 32 | 3274 | 0.0066866 | cellular_component | membrane |

**Turquoise module (2144 genes)**

Over-represented GO terms:

| category | num_in_subset | num_total | adj_pval | term | ontology |
|---|---|---|---|---|---|
| GO:0016021 | 811 | 2615 | 0.0000000 | cellular_component | integral component of membrane |
| GO:0016020 | 933 | 3274 | 0.0000000 | cellular_component | membrane |
| GO:0005887 | 215 | 435 | 0.0000000 | cellular_component | integral component of plasma membrane |
| GO:0005886 | 563 | 1833 | 0.0000000 | cellular_component | plasma membrane |
| GO:0005576 | 158 | 327 | 0.0000000 | cellular_component | extracellular region |
| GO:0005216 | 70 | 99 | 0.0000000 | molecular_function | ion channel activity |
| GO:0045202 | 117 | 251 | 0.0000000 | cellular_component | synapse |
| GO:0006811 | 113 | 244 | 0.0000000 | biological_process | ion transport |

| | | | | | |
|---|---|---|---|---|---|
| GO:0005615 | 147 | 384 | 0.0000000 | cellular_component | extracellular space |
| GO:0004930 | 77 | 150 | 0.0000000 | molecular_function | G protein-coupled receptor activity |
| GO:0007155 | 99 | 193 | 0.0000000 | biological_process | cell adhesion |
| GO:0007186 | 92 | 203 | 0.0000000 | biological_process | G protein-coupled receptor signaling pathway |
| GO:0007275 | 80 | 152 | 0.0000000 | biological_process | multicellular organism development |
| GO:0005509 | 144 | 361 | 0.0000000 | molecular_function | calcium ion binding |
| GO:0007156 | 47 | 65 | 0.0000000 | biological_process | homophilic cell adhesion via plasma membrane adhesion molecules |
| GO:0034220 | 50 | 83 | 0.0000000 | biological_process | ion transmembrane transport |
| GO:0007165 | 184 | 543 | 0.0000000 | biological_process | signal transduction |
| GO:0031012 | 59 | 95 | 0.0000000 | cellular_component | extracellular matrix |
| GO:0055085 | 127 | 329 | 0.0000000 | biological_process | transmembrane transport |
| GO:0007411 | 53 | 88 | 0.0000000 | biological_process | axon guidance |
| GO:0042391 | 37 | 52 | 0.0000000 | biological_process | regulation of membrane potential |
| GO:0005244 | 37 | 50 | 0.0000000 | molecular_function | voltage-gated ion channel activity |

| | | | | | |
|---|---|---|---|---|---|
| GO:0005604 | 38 | 48 | 0.0000000 | cellular_component | basement membrane |
| GO:0043005 | 69 | 142 | 0.0000000 | cellular_component | neuron projection |
| GO:0007268 | 41 | 65 | 0.0000000 | biological_process | chemical synaptic transmission |
| GO:0034765 | 37 | 52 | 0.0000000 | biological_process | regulation of ion transmembrane transport |
| GO:0030054 | 93 | 213 | 0.0000000 | cellular_component | cell junction |
| GO:0062023 | 37 | 52 | 0.0000000 | cellular_component | collagen-containing extracellular matrix |
| GO:0045211 | 37 | 57 | 0.0000000 | cellular_component | postsynaptic membrane |
| GO:0060078 | 21 | 23 | 0.0000000 | biological_process | regulation of postsynaptic membrane potential |
| GO:0098978 | 70 | 150 | 0.0000000 | cellular_component | glutamatergic synapse |
| GO:0099061 | 18 | 18 | 0.0000000 | cellular_component | integral component of postsynaptic density membrane |
| GO:1904315 | 19 | 20 | 0.0000000 | molecular_function | transmitter-gated ion channel activity involved in regulation of postsynaptic membrane potential |
| GO:0004714 | 26 | 31 | 0.0000000 | molecular_function | transmembrane receptor protein tyrosine kinase activity |
| GO:0030198 | 44 | 73 | 0.0000000 | biological_process | extracellular matrix organization |
| GO:0005201 | 25 | 28 | 0.0000000 | molecular_function | extracellular matrix structural constituent |

| | | | | | |
|---|---|---|---|---|---|
| GO:1902476 | 34 | 54 | 0.0000000 | biological_process | chloride transmembrane transport |
| GO:0043235 | 60 | 124 | 0.0000000 | cellular_component | receptor complex |
| GO:0004497 | 29 | 48 | 0.0000000 | molecular_function | monooxygenase activity |
| GO:0005254 | 23 | 32 | 0.0000000 | molecular_function | chloride channel activity |
| GO:0005230 | 16 | 19 | 0.0000001 | molecular_function | extracellular ligand-gated ion channel activity |
| GO:0005245 | 22 | 27 | 0.0000001 | molecular_function | voltage-gated calcium channel activity |
| GO:0050896 | 15 | 17 | 0.0000001 | biological_process | response to stimulus |
| GO:0006821 | 28 | 46 | 0.0000001 | biological_process | chloride transport |
| GO:0015276 | 15 | 16 | 0.0000002 | molecular_function | ligand-gated ion channel activity |
| GO:0030425 | 59 | 134 | 0.0000002 | cellular_component | dendrite |
| GO:0034707 | 17 | 21 | 0.0000002 | cellular_component | chloride channel complex |
| GO:0009986 | 89 | 249 | 0.0000002 | cellular_component | cell surface |
| GO:0005581 | 18 | 21 | 0.0000007 | cellular_component | collagen trimer |
| GO:0004890 | 11 | 11 | 0.0000009 | molecular_function | GABA-A receptor activity |
| GO:0008066 | 12 | 12 | 0.0000009 | molecular_function | glutamate receptor activity |

| | | | | | |
|---|---|---|---|---|---|
| GO:0030594 | 16 | 21 | 0.0000012 | molecular_function | neurotransmitter receptor activity |
| GO:0008395 | 15 | 19 | 0.0000013 | molecular_function | steroid hydroxylase activity |
| GO:0007605 | 38 | 70 | 0.0000014 | biological_process | sensory perception of sound |
| GO:0004970 | 12 | 12 | 0.0000022 | molecular_function | ionotropic glutamate receptor activity |
| GO:0070588 | 39 | 75 | 0.0000027 | biological_process | calcium ion transmembrane transport |
| GO:0035235 | 14 | 16 | 0.0000037 | biological_process | ionotropic glutamate receptor signaling pathway |
| GO:0005249 | 18 | 26 | 0.0000043 | molecular_function | voltage-gated potassium channel activity |
| GO:0007169 | 35 | 68 | 0.0000049 | biological_process | transmembrane receptor protein tyrosine kinase signaling pathway |
| GO:0016705 | 23 | 40 | 0.0000049 | molecular_function | oxidoreductase activity, acting on paired donors, with incorporation or reduction of molecular oxygen |
| GO:0004984 | 12 | 15 | 0.0000058 | molecular_function | olfactory receptor activity |
| GO:0050911 | 12 | 15 | 0.0000058 | biological_process | detection of chemical stimulus involved in sensory perception of smell |
| GO:0006816 | 37 | 72 | 0.0000086 | biological_process | calcium ion transport |

| | | | | | |
|---|---|---|---|---|---|
| GO:0030424 | 57 | 135 | 0.0000086 | cellular_component | axon |
| GO:0020037 | 30 | 62 | 0.0000132 | molecular_function | heme binding |
| GO:0045296 | 23 | 36 | 0.0000132 | molecular_function | cadherin binding |
| GO:0007399 | 47 | 109 | 0.0000142 | biological_process | nervous system development |
| GO:0004888 | 45 | 106 | 0.0000147 | molecular_function | transmembrane signaling receptor activity |
| GO:0016712 | 14 | 19 | 0.0000155 | molecular_function | oxidoreductase activity, acting on paired donors, with incorporation or reduction of molecular oxygen, reduced flavin or flavoprotein as one donor, and incorporation of one atom of oxygen |
| GO:0005891 | 15 | 18 | 0.0000185 | cellular_component | voltage-gated calcium channel complex |
| GO:0005267 | 21 | 35 | 0.0000191 | molecular_function | potassium channel activity |
| GO:0014069 | 42 | 93 | 0.0000259 | cellular_component | postsynaptic density |
| GO:0033674 | 25 | 42 | 0.0000259 | biological_process | positive regulation of kinase activity |
| GO:1902711 | 9 | 9 | 0.0000259 | cellular_component | GABA-A receptor complex |
| GO:0098609 | 40 | 83 | 0.0000274 | biological_process | cell-cell adhesion |

| | | | | | |
|---|---|---|---|---|---|
| GO:0031225 | 13 | 17 | 0.0000291 | cellular_component | anchored component of membrane |
| GO:0071805 | 32 | 70 | 0.0000372 | biological_process | potassium ion transmembrane transport |
| GO:0005262 | 27 | 47 | 0.0000436 | molecular_function | calcium channel activity |
| GO:0007214 | 13 | 17 | 0.0000649 | biological_process | gamma-aminobutyric acid signaling pathway |
| GO:0098982 | 17 | 24 | 0.0000649 | cellular_component | GABA-ergic synapse |
| GO:0008076 | 17 | 28 | 0.0000738 | cellular_component | voltage-gated potassium channel complex |
| GO:0005261 | 21 | 33 | 0.0000841 | molecular_function | cation channel activity |
| GO:0050804 | 27 | 52 | 0.0000912 | biological_process | modulation of chemical synaptic transmission |
| GO:0050877 | 14 | 20 | 0.0000912 | biological_process | nervous system process |
| GO:0004222 | 31 | 65 | 0.0001057 | molecular_function | metalloendopeptidase activity |
| GO:0038023 | 34 | 75 | 0.0001147 | molecular_function | signaling receptor activity |
| GO:0070509 | 14 | 17 | 0.0001147 | biological_process | calcium ion import |
| GO:0007218 | 15 | 23 | 0.0001194 | biological_process | neuropeptide signaling pathway |
| GO:0022851 | 8 | 8 | 0.0001704 | molecular_function | GABA-gated chloride ion channel activity |
| GO:0005506 | 31 | 73 | 0.0001749 | molecular_function | iron ion binding |

| | | | | | |
|---|---|---|---|---|---|
| GO:0051965 | 13 | 17 | 0.0001942 | biological_process | positive regulation of synapse assembly |
| GO:0035725 | 28 | 55 | 0.0001997 | biological_process | sodium ion transmembrane transport |
| GO:0030672 | 17 | 29 | 0.0002577 | cellular_component | synaptic vesicle membrane |
| GO:0008237 | 37 | 89 | 0.0002828 | molecular_function | metallopeptidase activity |
| GO:0008392 | 7 | 7 | 0.0003420 | molecular_function | arachidonic acid epoxygenase activity |
| GO:0019373 | 7 | 7 | 0.0003420 | biological_process | epoxygenase P450 pathway |
| GO:0005272 | 13 | 17 | 0.0004206 | molecular_function | sodium channel activity |
| GO:0008331 | 9 | 9 | 0.0004206 | molecular_function | high voltage-gated calcium channel activity |
| GO:0042734 | 14 | 21 | 0.0008151 | cellular_component | presynaptic membrane |
| GO:0043025 | 38 | 95 | 0.0008712 | cellular_component | neuronal cell body |
| GO:0006082 | 11 | 16 | 0.0009548 | biological_process | organic acid metabolic process |
| GO:0008083 | 20 | 42 | 0.0009691 | molecular_function | growth factor activity |
| GO:0032590 | 8 | 9 | 0.0009833 | cellular_component | dendrite membrane |
| GO:0031290 | 8 | 8 | 0.0010067 | biological_process | retinal ganglion cell axon guidance |
| GO:0098742 | 11 | 14 | 0.0011872 | biological_process | cell-cell adhesion via plasma-membrane adhesion molecules |

| | | | | | |
|---|---|---|---|---|---|
| GO:0022857 | 41 | 114 | 0.0012753 | molecular_function | transmembrane transporter activity |
| GO:0006508 | 85 | 289 | 0.0013567 | biological_process | proteolysis |
| GO:0005912 | 33 | 74 | 0.0013816 | cellular_component | adherens junction |
| GO:0086091 | 11 | 14 | 0.0014014 | biological_process | regulation of heart rate by cardiac conduction |
| GO:0006813 | 24 | 53 | 0.0018182 | biological_process | potassium ion transport |
| GO:0006814 | 22 | 45 | 0.0020032 | biological_process | sodium ion transport |
| GO:0043197 | 24 | 49 | 0.0021302 | cellular_component | dendritic spine |
| GO:0005003 | 8 | 9 | 0.0022232 | molecular_function | ephrin receptor activity |
| GO:0007216 | 8 | 9 | 0.0023318 | biological_process | G protein-coupled glutamate receptor signaling pathway |
| GO:0005005 | 8 | 9 | 0.0023757 | molecular_function | transmembrane-ephrin receptor activity |
| GO:0004867 | 16 | 29 | 0.0023759 | molecular_function | serine-type endopeptidase inhibitor activity |
| GO:0032281 | 10 | 14 | 0.0024915 | cellular_component | AMPA glutamate receptor complex |
| GO:0016339 | 11 | 15 | 0.0026130 | biological_process | calcium-dependent cell-cell adhesion via plasma membrane cell adhesion molecules |
| GO:0030317 | 19 | 36 | 0.0030522 | biological_process | flagellated sperm motility |

| | | | | | |
|---|---|---|---|---|---|
| GO:0005248 | 13 | 18 | 0.0033454 | molecular_function | voltage-gated sodium channel activity |
| GO:0099056 | 9 | 11 | 0.0038029 | cellular_component | integral component of presynaptic membrane |
| GO:0035249 | 11 | 16 | 0.0039103 | biological_process | synaptic transmission, glutamatergic |
| GO:0005518 | 16 | 29 | 0.0039662 | molecular_function | collagen binding |
| GO:0050808 | 16 | 28 | 0.0050015 | biological_process | synapse organization |
| GO:0015347 | 7 | 8 | 0.0055215 | molecular_function | sodium-independent organic anion transmembrane transporter activity |
| GO:0099560 | 10 | 14 | 0.0062109 | biological_process | synaptic membrane adhesion |
| GO:0001518 | 11 | 14 | 0.0065869 | cellular_component | voltage-gated sodium channel complex |
| GO:0016594 | 7 | 8 | 0.0071068 | molecular_function | glycine binding |
| GO:0007608 | 11 | 19 | 0.0085769 | biological_process | sensory perception of smell |
| GO:0008046 | 6 | 6 | 0.0098182 | molecular_function | axon guidance receptor activity |
| GO:0007160 | 22 | 46 | 0.0113008 | biological_process | cell-matrix adhesion |
| GO:2000311 | 7 | 8 | 0.0117409 | biological_process | regulation of AMPA receptor activity |
| GO:0008201 | 23 | 52 | 0.0118231 | molecular_function | heparin binding |

| | | | | | |
|---|---|---|---|---|---|
| GO:0005178 | 27 | 65 | 0.0122968 | molecular_function | integrin binding |
| GO:0009887 | 27 | 63 | 0.0133431 | biological_process | animal organ morphogenesis |
| GO:0004180 | 10 | 17 | 0.0136425 | molecular_function | carboxypeptidase activity |
| GO:0005237 | 5 | 5 | 0.0136425 | molecular_function | inhibitory extracellular ligand-gated ion channel activity |
| GO:0008503 | 5 | 5 | 0.0136425 | molecular_function | benzodiazepine receptor activity |
| GO:0060079 | 11 | 18 | 0.0136425 | biological_process | excitatory postsynaptic potential |
| GO:0034703 | 6 | 6 | 0.0136588 | cellular_component | cation channel complex |
| GO:0007631 | 5 | 5 | 0.0148946 | biological_process | feeding behavior |
| GO:0048013 | 12 | 21 | 0.0149937 | biological_process | ephrin receptor signaling pathway |
| GO:0060012 | 5 | 5 | 0.0156301 | biological_process | synaptic transmission, glycinergic |
| GO:0042738 | 9 | 15 | 0.0170549 | biological_process | exogenous drug catabolic process |
| GO:0045880 | 10 | 16 | 0.0175441 | biological_process | positive regulation of smoothened signaling pathway |
| GO:0034587 | 6 | 7 | 0.0191873 | biological_process | piRNA metabolic process |
| GO:0099060 | 6 | 7 | 0.0215254 | cellular_component | integral component of postsynaptic specialization membrane |
| GO:1904862 | 5 | 5 | 0.0215254 | biological_process | inhibitory synapse assembly |

| | | | | | |
|---|---|---|---|---|---|
| GO:0007601 | 20 | 47 | 0.0217298 | biological_process | visual perception |
| GO:0015721 | 8 | 11 | 0.0219669 | biological_process | bile acid and bile salt transport |
| GO:0042310 | 5 | 5 | 0.0219669 | biological_process | vasoconstriction |
| GO:0060077 | 5 | 5 | 0.0219669 | cellular_component | inhibitory synapse |
| GO:0016342 | 11 | 17 | 0.0225266 | cellular_component | catenin complex |
| GO:0019228 | 9 | 12 | 0.0225266 | biological_process | neuronal action potential |
| GO:0007269 | 9 | 14 | 0.0225356 | biological_process | neurotransmitter secretion |
| GO:0015106 | 7 | 9 | 0.0235010 | molecular_function | bicarbonate transmembrane transporter activity |
| GO:0030534 | 10 | 16 | 0.0235156 | biological_process | adult behavior |
| GO:0043252 | 6 | 7 | 0.0236664 | biological_process | sodium-independent organic anion transport |
| GO:0018108 | 31 | 80 | 0.0241308 | biological_process | peptidyl-tyrosine phosphorylation |
| GO:0007196 | 5 | 5 | 0.0256393 | biological_process | adenylate cyclase-inhibiting G protein-coupled glutamate receptor signaling pathway |
| GO:2000463 | 7 | 8 | 0.0256393 | biological_process | positive regulation of excitatory postsynaptic potential |

| | | | | | |
|---|---|---|---|---|---|
| GO:0007189 | 20 | 48 | 0.0269733 | biological_process | adenylate cyclase-activating G protein-coupled receptor signaling pathway |
| GO:0030414 | 10 | 16 | 0.0269733 | molecular_function | peptidase inhibitor activity |
| GO:0048018 | 6 | 8 | 0.0269733 | molecular_function | receptor ligand activity |
| GO:0045499 | 10 | 16 | 0.0280018 | molecular_function | chemorepellent activity |
| GO:0006805 | 12 | 25 | 0.0292781 | biological_process | xenobiotic metabolic process |
| GO:0007043 | 11 | 18 | 0.0292781 | biological_process | cell-cell junction assembly |
| GO:0016500 | 6 | 7 | 0.0310923 | molecular_function | protein-hormone receptor activity |
| GO:0042472 | 11 | 19 | 0.0323643 | biological_process | inner ear morphogenesis |
| GO:0022414 | 5 | 5 | 0.0328608 | biological_process | reproductive process |
| GO:0008528 | 9 | 15 | 0.0344847 | molecular_function | G protein-coupled peptide receptor activity |
| GO:0098685 | 18 | 40 | 0.0344847 | cellular_component | Schaffer collateral - CA1 synapse |
| GO:0017146 | 5 | 5 | 0.0358930 | cellular_component | NMDA selective glutamate receptor complex |
| GO:0086010 | 7 | 8 | 0.0362860 | biological_process | membrane depolarization during action potential |
| GO:0098793 | 25 | 64 | 0.0381558 | cellular_component | presynapse |

| | | | | | |
|---|---|---|---|---|---|
| GO:0030246 | 24 | 62 | 0.0392008 | molecular_function | carbohydrate binding |
| GO:0007626 | 17 | 36 | 0.0398328 | biological_process | locomotory behavior |
| GO:0016324 | 39 | 113 | 0.0398328 | cellular_component | apical plasma membrane |
| GO:0051015 | 41 | 116 | 0.0421992 | molecular_function | actin filament binding |
| GO:0017046 | 8 | 12 | 0.0432277 | molecular_function | peptide hormone binding |
| GO:0010466 | 10 | 17 | 0.0443575 | biological_process | negative regulation of peptidase activity |
| GO:0007158 | 6 | 7 | 0.0454588 | biological_process | neuron cell-cell adhesion |
| GO:0030199 | 11 | 19 | 0.0454588 | biological_process | collagen fibril organization |
| GO:0042626 | 21 | 47 | 0.0454588 | molecular_function | ATPase-coupled transmembrane transporter activity |
| GO:0098839 | 8 | 12 | 0.0454588 | cellular_component | postsynaptic density membrane |
| GO:0003779 | 63 | 199 | 0.0466297 | molecular_function | actin binding |
| GO:0044458 | 8 | 12 | 0.0466297 | biological_process | motile cilium assembly |
| GO:0098688 | 6 | 7 | 0.0466297 | cellular_component | parallel fiber to Purkinje cell synapse |
| GO:0036477 | 7 | 10 | 0.0467352 | cellular_component | somatodendritic compartment |
| GO:0003774 | 33 | 79 | 0.0471490 | molecular_function | motor activity |

| | | | | | |
|---|---|---|---|---|---|
| GO:0050649 | 4 | 4 | 0.0471490 | molecular_function | testosterone 6-beta-hydroxylase activity |
| GO:0050919 | 12 | 22 | 0.0488865 | biological_process | negative chemotaxis |

Under-represented GO terms:

| category | num_in_subset | num_total | adj_pval | term | ontology |
|---|---|---|---|---|---|
| GO:0003723 | 42 | 519 | 0.0000000 | molecular_function | RNA binding |
| GO:0005634 | 287 | 3117 | 0.0000000 | cellular_component | nucleus |
| GO:0005654 | 180 | 2064 | 0.0000000 | cellular_component | nucleoplasm |
| GO:0005730 | 24 | 501 | 0.0000000 | cellular_component | nucleolus |
| GO:0005739 | 47 | 803 | 0.0000000 | cellular_component | mitochondrion |
| GO:0005813 | 25 | 360 | 0.0000000 | cellular_component | centrosome |
| GO:0005829 | 215 | 2063 | 0.0000000 | cellular_component | cytosol |
| GO:0006974 | 13 | 247 | 0.0000000 | biological_process | cellular response to DNA damage stimulus |
| GO:0016607 | 13 | 262 | 0.0000000 | cellular_component | nuclear speck |
| GO:0006412 | 2 | 164 | 0.0000001 | biological_process | translation |

| | | | | | |
|---|---|---|---|---|---|
| GO:0016604 | 12 | 229 | 0.0000001 | cellular_component | nuclear body |
| GO:0006511 | 6 | 167 | 0.0000003 | biological_process | ubiquitin-dependent protein catabolic process |
| GO:0006325 | 1 | 103 | 0.0000003 | biological_process | chromatin organization |
| GO:0005737 | 438 | 2821 | 0.0000021 | cellular_component | cytoplasm |
| GO:0031625 | 8 | 177 | 0.0000159 | molecular_function | ubiquitin protein ligase binding |
| GO:0016567 | 25 | 311 | 0.0000161 | biological_process | protein ubiquitination |
| GO:0008168 | 1 | 99 | 0.0000194 | molecular_function | methyltransferase activity |
| GO:0000398 | 2 | 108 | 0.0000957 | biological_process | mRNA splicing, via spliceosome |
| GO:0032259 | 1 | 90 | 0.0000957 | biological_process | methylation |
| GO:0045892 | 27 | 301 | 0.0000957 | biological_process | negative regulation of transcription, DNA-templated |
| GO:0006281 | 9 | 159 | 0.0001723 | biological_process | DNA repair |
| GO:0004386 | 2 | 86 | 0.0002110 | molecular_function | helicase activity |
| GO:0000122 | 49 | 448 | 0.0004651 | biological_process | negative regulation of transcription by RNA polymerase II |
| GO:0003735 | 1 | 104 | 0.0004719 | molecular_function | structural constituent of ribosome |
| GO:0005764 | 9 | 161 | 0.0007528 | cellular_component | lysosome |

| | | | | | |
|---|---|---|---|---|---|
| GO:0016740 | 91 | 705 | 0.0007799 | molecular_function | transferase activity |
| GO:0000785 | 51 | 455 | 0.0008320 | cellular_component | chromatin |
| GO:0005681 | 0 | 69 | 0.0008661 | cellular_component | spliceosomal complex |
| GO:0015031 | 15 | 204 | 0.0008661 | biological_process | protein transport |
| GO:0005840 | 1 | 99 | 0.0008752 | cellular_component | ribosome |
| GO:0018105 | 5 | 111 | 0.0009947 | biological_process | peptidyl-serine phosphorylation |
| GO:0071013 | 0 | 62 | 0.0012557 | cellular_component | catalytic step 2 spliceosome |
| GO:0042393 | 6 | 114 | 0.0016338 | molecular_function | histone binding |
| GO:0003682 | 29 | 285 | 0.0016598 | molecular_function | chromatin binding |
| GO:0004842 | 14 | 179 | 0.0023393 | molecular_function | ubiquitin-protein transferase activity |
| GO:0003713 | 11 | 151 | 0.0024953 | molecular_function | transcription coactivator activity |
| GO:0005525 | 16 | 216 | 0.0024953 | molecular_function | GTP binding |
| GO:0003676 | 47 | 418 | 0.0032570 | molecular_function | nucleic acid binding |
| GO:0003677 | 96 | 698 | 0.0056666 | molecular_function | DNA binding |
| GO:0016579 | 1 | 59 | 0.0058472 | biological_process | protein deubiquitination |

| | | | | | |
|---|---|---|---|---|---|
| GO:0035861 | 0 | 50 | 0.0065521 | cellular_component | site of double-strand break |
| GO:0042826 | 2 | 70 | 0.0077110 | molecular_function | histone deacetylase binding |
| GO:0019899 | 18 | 200 | 0.0094879 | molecular_function | enzyme binding |
| GO:0035064 | 0 | 46 | 0.0094879 | molecular_function | methylated histone binding |
| GO:0006355 | 83 | 629 | 0.0112499 | biological_process | regulation of transcription, DNA-templated |
| GO:0032991 | 38 | 333 | 0.0165935 | cellular_component | protein-containing complex |
| GO:0000724 | 1 | 55 | 0.0192048 | biological_process | double-strand break repair via homologous recombination |
| GO:0003678 | 0 | 39 | 0.0196822 | molecular_function | DNA helicase activity |
| GO:0008380 | 3 | 83 | 0.0227186 | biological_process | RNA splicing |
| GO:0006888 | 1 | 59 | 0.0246812 | biological_process | endoplasmic reticulum to Golgi vesicle-mediated transport |
| GO:0006886 | 17 | 188 | 0.0283941 | biological_process | intracellular protein transport |
| GO:0006357 | 99 | 712 | 0.0296350 | biological_process | regulation of transcription by RNA polymerase II |
| GO:0032508 | 1 | 49 | 0.0296350 | biological_process | DNA duplex unwinding |
| GO:0000932 | 1 | 53 | 0.0306984 | cellular_component | P-body |
| GO:0003725 | 1 | 53 | 0.0306984 | molecular_function | double-stranded RNA binding |

| | | | | | |
|---|---|---|---|---|---|
| GO:0006397 | 8 | 124 | 0.0306984 | biological_process | mRNA processing |
| GO:0005635 | 5 | 88 | 0.0352408 | cellular_component | nuclear envelope |
| GO:0045944 | 89 | 627 | 0.0352408 | biological_process | positive regulation of transcription by RNA polymerase II |
| GO:0045893 | 49 | 380 | 0.0381968 | biological_process | positive regulation of transcription, DNA-templated |
| GO:0006364 | 1 | 54 | 0.0439499 | biological_process | rRNA processing |

**Orange module (47 genes)**

Over-represented GO terms:

| category | num_in_subset | num_total | adj_pval | term | ontology |
|---|---|---|---|---|---|
| GO:0005884 | 6 | 44 | 0.0166133 | cellular_component | actin filament |

**Green module (220 genes)**

Over-represented GO terms:

| category | num_in_subset | num_total | adj_pval | term | ontology |
|---|---|---|---|---|---|
| GO:0022625 | 11 | 33 | 0.0248793 | cellular_component | cytosolic large ribosomal subunit |

**Tan module (194 genes)**

Under-represented GO terms:

| category | num_in_subset | num_total | adj_pval | term | ontology |
|---|---|---|---|---|---|
| GO:0016021 | 27 | 2615 | 0.0307406 | cellular_component | integral component of membrane |

**Light green module (91 genes)**

Over-represented GO terms:

| category | num_in_subset | num_total | adj_pval | term | ontology |
|---|---|---|---|---|---|

| | | | | | |
|---|---|---|---|---|---|
| GO:0035914 | 6 | 28 | 0.0370413 | biological_process | skeletal muscle cell differentiation |
