## Supplementary material for "Inference of Rift Valley Fever pathogenesis in *Bos taurus* using a gene co-expression network": non-redundant biological processes GO terms in the turquoise module

**Additional file 5: Table S3.** A summary of non-redundant biological processes GO terms in the turquoise module.

| **Term_ID** | **Name** |
|---|---|
| GO:0007605 | sensory perception of sound |
| GO:0007631 | feeding behavior |
| GO:0030534 | adult behavior |
| GO:0007626 | locomotory behavior |
| GO:0042391 | regulation of membrane potential |
| GO:0050896 | response to stimulus |
| GO:1902476 | chloride transmembrane transport |
| GO:0043252 | sodium-independent organic anion transport |
| GO:0015721 | bile acid and bile salt transport |
| GO:0070588 | calcium ion transmembrane transport |
| GO:0006813 | potassium ion transport |
| GO:0006814 | sodium ion transport |
| GO:0034220 | ion transmembrane transport |
| GO:0071805 | potassium ion transmembrane transport |
| GO:0035725 | sodium ion transmembrane transport |
| GO:0006816 | calcium ion transport |
| GO:0070509 | calcium ion import |
| GO:0006821 | chloride transport |
| GO:0019373 | epoxygenase P450 pathway |
| GO:0030198 | extracellular matrix organization |
| GO:0030199 | collagen fibril organization |
| GO:0007268 | chemical synaptic transmission |

| | |
|---|---|
| GO:0007214 | gamma-aminobutyric acid signaling pathway |
| GO:0060012 | synaptic transmission |
| GO:0035249 | synaptic transmission |
| GO:0060079 | excitatory postsynaptic potential |
| GO:0007269 | neurotransmitter secretion |
| GO:0007155 | cell adhesion |
| GO:0007158 | neuron cell-cell adhesion |
| GO:0007160 | cell-matrix adhesion |
| GO:0007156 | homophilic cell adhesion via plasma membrane adhesion molecules |
| GO:0099560 | synaptic membrane adhesion |
| GO:0016339 | calcium-dependent cell-cell adhesion via plasma membrane cell adhesion molecules |
| GO:0098742 | cell-cell adhesion via plasma-membrane adhesion molecules |
| GO:0098609 | cell-cell adhesion |
| GO:0034765 | regulation of ion transmembrane transport |
| GO:2000311 | regulation of AMPA receptor activity |
| GO:0050804 | modulation of chemical synaptic transmission |
| GO:2000463 | positive regulation of excitatory postsynaptic potential |
| GO:0033674 | positive regulation of kinase activity |
| GO:0034587 | piRNA metabolic process |
| GO:0018108 | peptidyl-tyrosine phosphorylation |
| GO:0035235 | ionotropic glutamate receptor signaling pathway |
| GO:0007196 | adenylate cyclase-inhibiting G protein-coupled glutamate receptor signaling pathway |
| GO:0007216 | G protein-coupled glutamate receptor signaling pathway |

| | |
|---|---|
| GO:0007218 | neuropeptide signaling pathway |
| GO:0007411 | axon guidance |
| GO:0044458 | motile cilium assembly |
| GO:0030317 | flagellated sperm motility |
| GO:0009887 | animal organ morphogenesis |
| GO:0022414 | reproductive process |
| GO:0050919 | negative chemotaxis |
| GO:0031290 | retinal ganglion cell axon guidance |
| GO:0050808 | synapse organization |
| GO:1904862 | inhibitory synapse assembly |
| GO:0007043 | cell-cell junction assembly |
| GO:0042738 | exogenous drug catabolic process |
| GO:0006805 | xenobiotic metabolic process |
| GO:0051965 | positive regulation of synapse assembly |
| GO:0060078 | regulation of postsynaptic membrane potential |
| GO:0006508 | proteolysis |
| GO:0086091 | regulation of heart rate by cardiac conduction |
| GO:0019228 | neuronal action potential |
| GO:0086010 | membrane depolarization during action potential |
| GO:0045880 | positive regulation of smoothened signaling pathway |
| GO:0055085 | transmembrane transport |
| GO:0006811 | ion transport |
| GO:0042310 | vasoconstriction |
| GO:0007165 | signal transduction |
| GO:0007186 | G protein-coupled receptor signaling pathway |

| | |
|---|---|
| GO:0007275 | multicellular organism development |
| GO:0007399 | nervous system development |
| GO:0007169 | transmembrane receptor protein tyrosine kinase signaling pathway |
| GO:0048013 | ephrin receptor signaling pathway |
| GO:0007189 | adenylate cyclase-activating G protein-coupled receptor signaling pathway |
| GO:0042472 | inner ear morphogenesis |
| GO:0006082 | organic acid metabolic process |
| GO:0007601 | visual perception |
| GO:0050911 | detection of chemical stimulus involved in sensory perception of smell |
| GO:0050877 | nervous system process |
| GO:0007608 | sensory perception of smell |
| GO:0010466 | negative regulation of peptidase activity |
