## Supplementary material for "Inference of Rift Valley Fever pathogenesis in *Bos taurus* using a gene co-expression network": non-redundant molecular function GO terms in the turquoise module

**Additional file 6: Table S4.** A summary of the non-redundant molecular function GO terms in the turquoise module.

| Term_ID | Name |
|---|---|
| GO:0004497 | monooxygenase activity |
| GO:0005201 | extracellular matrix structural constituent |
| GO:0005509 | calcium ion binding |
| GO:1904315 | transmitter-gated ion channel activity involved in regulation of postsynaptic membrane potential |
| GO:0005244 | voltage-gated ion channel activity |
| GO:0008503 | benzodiazepine receptor activity |
| GO:0008331 | high voltage-gated calcium channel activity |
| GO:0005237 | inhibitory extracellular ligand-gated ion channel activity |
| GO:0005216 | ion channel activity |
| GO:0005245 | voltage-gated calcium channel activity |
| GO:0004970 | ionotropic glutamate receptor activity |
| GO:0005262 | calcium channel activity |
| GO:0015276 | ligand-gated ion channel activity |
| GO:0005261 | cation channel activity |
| GO:0005249 | voltage-gated potassium channel activity |
| GO:0005267 | potassium channel activity |
| GO:0005230 | extracellular ligand-gated ion channel activity |
| GO:0017046 | peptide hormone binding |
| GO:0005518 | collagen binding |
| GO:0045296 | cadherin binding |
| GO:0005178 | integrin binding |

| | |
|---|---|
| GO:0008201 | heparin binding |
| GO:0020037 | heme binding |
| GO:0030246 | carbohydrate binding |
| GO:0004867 | serine-type endopeptidase inhibitor activity |
| GO:0030414 | peptidase inhibitor activity |
| GO:0003774 | motor activity |
| GO:0003779 | actin binding |
| GO:0008083 | growth factor activity |
| GO:0048018 | receptor ligand activity |
| GO:0016594 | glycine binding |
| GO:0008046 | axon guidance receptor activity |
| GO:0016500 | protein-hormone receptor activity |
| GO:0004890 | GABA-A receptor activity |
| GO:0022851 | GABA-gated chloride ion channel activity |
| GO:0005506 | iron ion binding |
| GO:0004714 | transmembrane receptor protein tyrosine kinase activity |
| GO:0005005 | transmembrane-ephrin receptor activity |
| GO:0005003 | ephrin receptor activity |
| GO:0004222 | metalloendopeptidase activity |
| GO:0008066 | glutamate receptor activity |
| GO:0030594 | neurotransmitter receptor activity |
| GO:0008528 | G protein-coupled peptide receptor activity |
| GO:0042626 | ATPase-coupled transmembrane transporter activity |
| GO:0008395 | steroid hydroxylase activity |
| GO:0008392 | arachidonic acid epoxygenase activity |

| | |
|---|---|
| GO:0016712 | oxidoreductase activity |
| GO:0004180 | carboxypeptidase activity |
| GO:0004930 | G protein-coupled receptor activity |
| GO:0004984 | olfactory receptor activity |
| GO:0004888 | transmembrane signaling receptor activity |
| GO:0038023 | signaling receptor activity |
| GO:0005272 | sodium channel activity |
| GO:0005248 | voltage-gated sodium channel activity |
| GO:0045499 | chemorepellent activity |
| GO:0016705 | oxidoreductase activity |
| GO:0005254 | chloride channel activity |
| GO:0015347 | sodium-independent organic anion transmembrane transporter activity |
| GO:0008237 | metallopeptidase activity |
| GO:0015106 | bicarbonate transmembrane transporter activity |
| GO:0022857 | transmembrane transporter activity |
| GO:0050649 | testosterone 6-beta-hydroxylase activity |
| GO:0051015 | actin filament binding |
