## Supplementary material for "Inference of Rift Valley Fever pathogenesis in *Bos taurus* using a gene co-expression network": non-redundant cellular compartment GO terms in the turquoise module

**Additional file 7: Table S5.** A summary of the non-redundant cellular compartment GO terms in the turquoise module.

| **Term_ID** | **Name** |
|---|---|
| GO:0043235 | receptor complex |
| GO:0045202 | synapse |
| GO:0031225 | anchored component of membrane |
| GO:0031012 | extracellular matrix |
| GO:0005604 | basement membrane |
| GO:0062023 | collagen-containing extracellular matrix |
| GO:0036477 | somatodendritic compartment |
| GO:0009986 | cell surface |
| GO:0043005 | neuron projection |
| GO:0030054 | cell junction |
| GO:0005615 | extracellular space |
| GO:0016020 | membrane |
| GO:0005576 | extracellular region |
| GO:1902711 | GABA-A receptor complex |
| GO:0034707 | chloride channel complex |
| GO:0005581 | collagen trimer |
| GO:0016021 | integral component of membrane |
| GO:0005886 | plasma membrane |
| GO:0016342 | catenin complex |
| GO:0005887 | integral component of plasma membrane |
| GO:0060077 | inhibitory synapse |
| GO:0098688 | parallel fiber to Purkinje cell synapse |

| | |
|---|---|
| GO:0098982 | GABA-ergic synapse |
| GO:0098685 | Schaffer collateral - CA1 synapse |
| GO:0030672 | synaptic vesicle membrane |
| GO:0005912 | adherens junction |
| GO:0045211 | postsynaptic membrane |
| GO:0099061 | integral component of postsynaptic density membrane |
| GO:0014069 | postsynaptic density |
| GO:0099056 | integral component of presynaptic membrane |
| GO:0042734 | presynaptic membrane |
| GO:0098839 | postsynaptic density membrane |
| GO:0099060 | integral component of postsynaptic specialization membrane |
| GO:0098978 | glutamatergic synapse |
| GO:0016324 | apical plasma membrane |
| GO:0030425 | dendrite |
| GO:0030424 | axon |
| GO:0032590 | dendrite membrane |
| GO:0043197 | dendritic spine |
| GO:0043025 | neuronal cell body |
| GO:0001518 | voltage-gated sodium channel complex |
| GO:0098793 | presynapse |
| GO:0005891 | voltage-gated calcium channel complex |
| GO:0032281 | AMPA glutamate receptor complex |
| GO:0008076 | voltage-gated potassium channel complex |
| GO:0034703 | cation channel complex |

| | |
|---|---|
| GO:0017146 | NMDA selective glutamate receptor complex |
