## Supplementary material for "Inference of Rift Valley Fever pathogenesis in *Bos taurus* using a gene co-expression network": non-redundant biological function GO terms in the red module

**Additional file 8: Table S6.** A summary of the non-redundant biological function GO terms in the red module.

| **Term_ID** | **Description** |
|---|---|
| GO:0016567 | protein ubiquitination |
| GO:0033173 | calcineurin-NFAT signalling cascade |
| GO:0006511 | ubiquitin-dependent protein catabolic process |
