## Supplementary material for "Inference of Rift Valley Fever pathogenesis in *Bos taurus* using a gene co-expression network": non-redundant molecular function GO terms in the red module

**Additional file 10: Table S8.** A summary of the non-redundant molecular function GO terms in the red module.

| **Term_ID** | **Name** |
| --- | --- |
| GO:0003677 | DNA binding |
| GO:0004842 | ubiquitin-protein transferase activity |
| GO:0061630 | ubiquitin protein ligase activity |
