## Supplementary material for "Inference of Rift Valley Fever pathogenesis in *Bos taurus* using a gene co-expression network": non-redundant Cellular compartment Gene Ontology terms in the red module

**Additional file 10: Table S8.** A summary of non-redundant Cellular compartment Gene Ontology terms in the red module.

| **Term_ID** | **Name** |
|---|---|
| GO:0005634 | nucleus |
| GO:0005829 | cytosol |
| GO:0005654 | nucleoplasm |
