## Supplementary material for "Inference of Rift Valley Fever pathogenesis in *Bos taurus* using a gene co-expression network": non-redundant cellular compartment GO terms in the white module

**Additional file 13: Table S11.** A summary of non-redundant cellular compartment GO terms in the white module.

| Term_ID | Name |
|---|---|
| GO:0005840 | ribosome |
| GO:0022627 | cytosolic small ribosomal subunit |
