## Supplementary material for "Inference of Rift Valley Fever pathogenesis in *Bos taurus* using a gene co-expression network": non-redundant Molecular function GO terms in the Steelblue module

**Additional file 15: Table S13.** A summary of non-redundant Molecular function GO terms in the Steelblue module.

| **Term_ID** | **Name** |
|---|---|
| GO:0003735 | structural constituent of ribosome |
