## Supplementary material for "Inference of Rift Valley Fever pathogenesis in *Bos taurus* using a gene co-expression network": Cellular compartment GO terms in the cyan module

**Additional file 21: Table S19.** A summary of non-redundant Cellular compartment GO terms in the cyan module.

| **Term_ID** | **Name** |
|---|---|
| GO:0016020 | membrane |
| GO:0016021 | integral component of membrane |
